# Resistance to Pyrethroids in *Aedes aegypti*: Insights into Transcriptomic Response to Different Insecticide Concentrations Transcriptomic responses of *Aedes aegypti* to insecticide concentrations

**DOI:** 10.64898/2026.03.12.711295

**Authors:** Alejandro Mejía Muñoz, Ana María Mejía-Jaramillo, Carl Lowenberger, Karla Saavedra Rodríguez, Omar Triana-Chávez

## Abstract

Insecticide spraying is a common strategy for controlling dengue outbreaks, but its effectiveness is compromised by the development of resistance in mosquito populations.

In this study, we subjected a strain of *Aedes aegypti* known for its exceptional ability to develop resistance to controlled permethrin and lambda-cyhalothrin insecticides pressure using two different concentrations. We analyzed resistance mechanisms that are enhanced at each concentration and used RNA sequencing to identify transcripts specifically associated with these exposure levels. Our objective was to uncover the molecular mechanisms triggered by different insecticide concentrations and to distinguish responses between type I and type II pyrethroids, which differ in chemical structure. Our results showed that *kdr* mutations confer only moderate levels of resistance, as do detoxifying enzymes.

For lambda-cyhalothrin, we identified genes involved in the electron transport chain, mitochondrial function, and overall responses to oxidative stress. tRNA transcripts were also upregulated, along with mitochondrial and stress-response transcripts, suggesting a metabolic shift, particularly toward maintaining homeostasis under oxidative stress. These changes point to mechanisms that sustain resistance to this type II insecticide beyond direct detoxification in this population.

On the contrary, permethrin induced marked overexpression of cuticle genes, CYP450 genes (especially CYP4), and Odorant Binding Proteins. These expression patterns, together with metabolic enzymes, point to detoxification, reduced penetration, or even sequestration of insecticide, all of which intensify with increasing concentrations. This overregulation of genes suggests an integrated response complemented by classical metabolic detoxification and accompanied by overregulation of mitochondrial complexes.

We showed that despite the shared mode of action of the insecticides permethrin and lambda-cyhalothrin, they elicit distinct responses in this *Ae. aegypti* population. We also showed that the transcriptomic response depends on insecticide concentration and may modulate insecticide tolerance. This article advances understanding of the complexity of pyrethroid resistance in *Aedes aegypti* and underscores the importance of considering both the insecticide type and the concentration used in vector control programs.

**Author summary:** *Aedes aegypti* mosquitoes transmit dengue and other arboviruses, being a major public health problem in tropical regions like Colombia, where control relies on pyrethroid insecticide spraying. Based on reports of inconsistent results in the field due to different effects of insecticide concentrations, we recreated variable doses by exposing a resistant Colombian *Aedes aegypti* strain to low (LC25) and high (LC75) concentrations of permethrin (type I) and lambda-cyhalothrin (type II) to identify concentration-dependent resistance mechanisms. Using genetic mutation analysis, enzyme activity assays, and RNA sequencing, we identified the molecular mechanisms these mosquitoes use to survive.

Knockdown resistance (kdr) and detoxification enzymes contributed to some extent to resistance but varied by insecticide type and concentration. RNAseq identified that lambda-cyhalothrin upregulated genes for mitochondrial energy production, oxidative stress defense, immune signaling, and transfer RNAs, facilitating homeostasis under chemical stressors. Permethrin instead upregulated genes for cuticle thickening, cytochrome P450 enzymes, and odorant-binding proteins, which are associated with improved penetration barriers, and metabolic breakdown that intensified with higher concentrations.

This reveals pyrethroid resistance as complex beyond classic mechanisms, as even low field doses favor stress tolerance or physical defenses to evade sprays. We detected transcripts that improve survival at high concentrations and could be selected in these mosquitoes. Carefully selecting the type of pyrethroid to be used and the dose should be an important factor in vector control. This optimizes current interventions, prolongs their efficacy, and aids researchers in modeling resistance to protect communities.

## Introduction

Pyrethroid insecticides are commonly used to control *Aedes aegypti* because they cause fewer nuisance and adverse effects in humans than organophosphates [1,2]. However, widespread and continued use of pyrethroids has led to the selection of resistance in mosquito populations [3,4]. Currently, the main resistance mechanisms include *kdr* mutations in the voltage-gated sodium channel (*vgsc*) gene [5] and metabolic resistance mediated by detoxification enzymes that modify or break down the insecticide [6]. Additional mechanisms have also been reported, highlighting the complexity of insecticide resistance (IR) and the challenge of predicting its outcomes [7].

Despite their widespread use, translating insecticide applications into effective vector control has proven challenging. Field interventions often yield inconsistent results [8,9], largely due to inadequate quality control and incomplete household coverage [8,10]. Furthermore, spraying does not reliably deliver lethal doses throughout the household environment [10], and the repellent effect of pyrethroids is short-lived [11].

While new insecticide molecules and formulations are being explored, their discovery and approval are both expensive and time-intensive. In the meantime, identifying and characterizing resistance mechanisms is essential for sustaining the effectiveness of existing tools and optimizing control strategies.

Understanding how varying insecticide concentrations affect mosquitoes is essential for resistance management [12]. Exposing insects to both lethal and sublethal concentrations provides insight into the spectrum of resistance and improves allele detection by decreasing false positives [13]. In general, low concentrations select for cumulative traits with synergistic effects, while high concentrations favor rare mutations with strong resistance phenotypes [12].

This study aimed to elucidate how resistance mechanisms differ between type I (permethrin) and type II (lambda-cyhalothrin) pyrethroids and across varying insecticide concentrations. To address this, we examined three key components of resistance: the frequency of kdr alleles in the voltage-gated sodium channel gene, the activity of detoxification enzymes, and transcriptomic responses using RNA sequencing (RNA-seq). Our general hypothesis is that resistance mechanisms in *Ae. aegypti* differ depending on both the pyrethroid class (type I vs. type II) and the exposure concentration, with distinct contributions from target-site mutations, metabolic detoxification, and gene expression responses. Our results have allowed us to identify additional mechanisms of insecticide resistance by RNA-seq, which vary depending on the type and concentration of pyrethroid used, mainly in the form of responses to oxidative stress and integrated mechanisms of resistance that have the potential to slow down damage to the molecular target, not only based on kdr mutations or metabolic detoxification.

## Materials and Methods

### Mosquitoes

The *Ae. aegypti* population used in this study was originally collected in Acacías, Meta, Colombia, in 2016 [14] and designated as the AF strain. Mosquito colonies were maintained under controlled insectary conditions (25 ± 5 °C, 72% ± 5% relative humidity, and a 12:12h light:dark photoperiod). Larvae were reared in dechlorinated water and fed Purina® Truchina fish food (48% protein). Upon emergence, adults were transferred to 50 × 50 × 50 cm cages and supplied with a 10% sugar solution ad libitum. Females were blood-fed on mice, and oviposition substrates consisted of a strip of moistened filter paper. The egg strips were collected and stored in humidity-controlled boxes for later use. The AF population was maintained for 13 generations without insecticide exposure. Each generation consisted of 1,000 mosquitoes of both sexes. Despite the absence of selective pressure, this colony preserved its innate resistance profile [15].

### Experimental design and pyrethroid selection procedure

To assess resistance under insecticide pressure, a subset of the AF population was exposed for two successive generations to either a high (H) or a low (L) concentration of permethrin (P) or lambda-cyhalothrin (L). Selection pressure bioassays were conducted using the Centers for Disease Control and Prevention (CDC) bottle bioassays (Fig 1). First, we tested different insecticide concentrations to estimate the diagnostic dose (DD) and diagnostic time (DT) for each insecticide in AF adults, as explained below. Using these data, we calculated the concentrations that killed 75% and 25% (LC_75_ and LC_25_, respectively). To confirm resistance, we conducted larval bioassays following World Health Organization (WHO) protocols [16] in the original AF population, the pyrethroid-selected strains, and the Rockefeller strain, a susceptible reference strain. Resistance ratios (RR) were calculated as the LC_50_ ratio between the tested population and the susceptible reference population. Resistance criteria were categorized as follows: a value above 10-fold indicates resistance, a value between 5-and 10-fold indicates moderate resistance, and a value below 5-fold indicates susceptibility [17].

**Fig 1.**
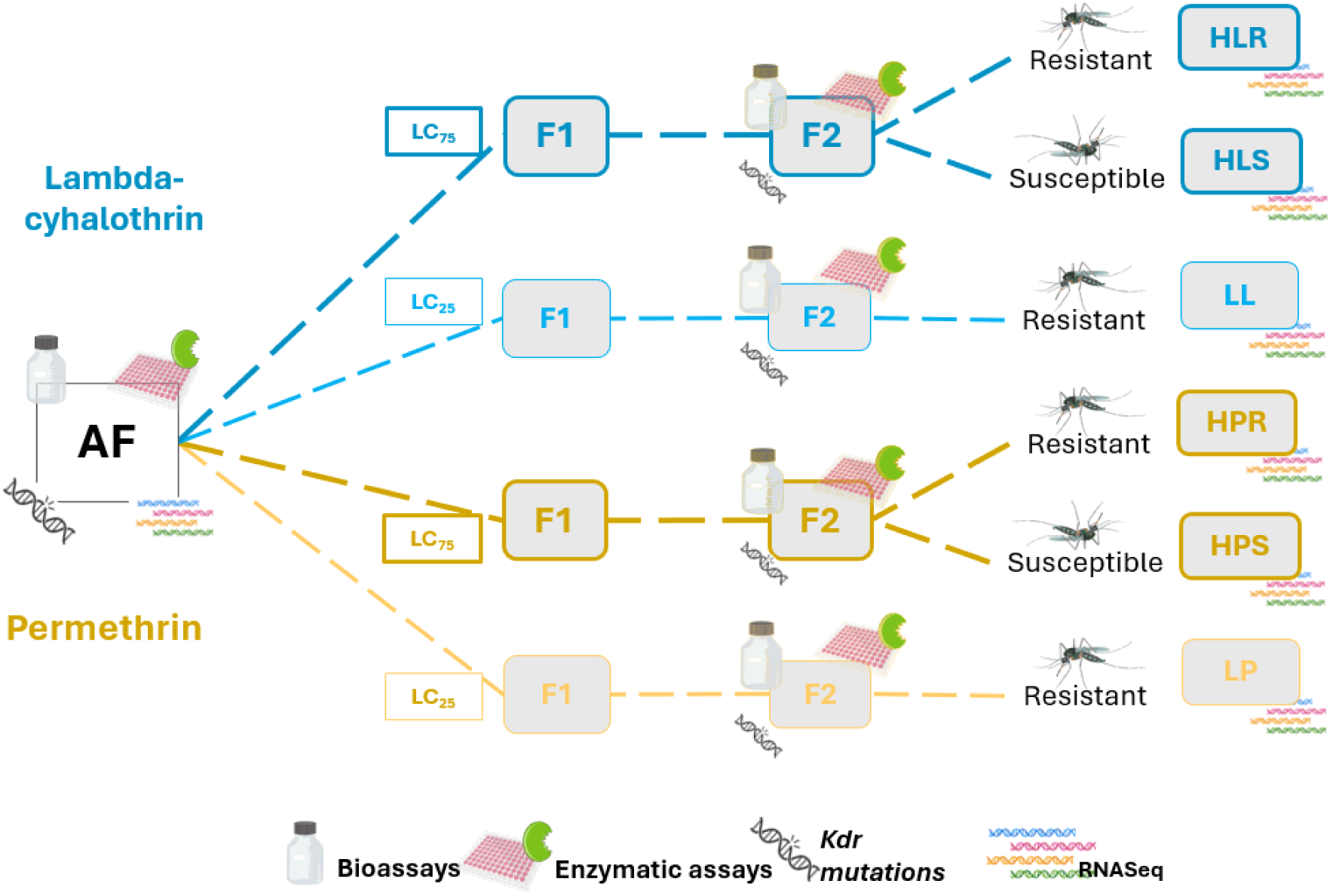
Overview of the experimental design. Exposure to insecticides is represented by dashed colored lines, with each lethal concentration (LC) depicted in colored boxes. Different LC pressures are also indicated by dark blue (AF population exposed to Lambda-cyhalothrin at LC_75_), light blue (AF population exposed to lambda-cyhalothrin at LC_25_), dark yellow (AF population exposed to Permethrin at LC_75_), and light yellow (AF population exposed to Permethrin at LC_25_). We used CDC adult bioassays to calculate DD and DT and to conduct the selection pressure. DD and DT represent the concentration and exposure time, respectively, needed to achieve total knockdown of mosquitoes within a population. Mortality was measured based on these DD and DT in AF, F1, and F2 for each insecticide pressure line. WHO larvae bioassays were used to determine resistance ratios (RR) in AF and F2 for each insecticide pressure line. Enzymatic assays and *kdr* allele phenotyping were carried out on AF and F2 of each strain, and RNA-seq analysis was performed on AF and on resistant and susceptible populations of the F2 generation.

### Selection pressure bioassays: Diagnostic dose (DD) and diagnostic time (DT)

To establish selective pressure conditions for the AF strain, we first determined the diagnostic dose (DD) and diagnostic time (DT). Bottle bioassays were carried out according to CDC protocols [18] using 2-to 5-day-old female mosquitoes. We used lambda-cyhalothrin (98.7% a.i., Lot SZBB332XV, Sigma-Aldrich; Buchs, Switzerland), permethrin (98.9% active ingredient [a.i.], Lot 7042300, Chem Service; West Chester, Pennsylvania, USA), and, as a control, molecular-grade ethanol (Pan Reac Applichem; Lot 0V011514; Darmstadt, Germany). For each insecticide, three replicates of 20 female mosquitoes were exposed to five concentrations ranging from 6 to 72 ppm for lambda-cyhalothrin and from 30 to 300 ppm for permethrin. Following CDC criteria [18], mosquitoes exhibiting “knockdown” behavior were classified as dead. However, because some individuals may recover within 24 hours [19], we adopted the modification proposed by da-Cunha et al. (2005), scoring only the absence of movement as knockdown [20]. Mortality was recorded every 5 minutes for 1 hour, and DD and DT were determined as the times at which 100% of mosquitoes experienced knockdown.

### Selection pressure bioassays: Lethal concentration

To establish low and high concentrations for our selection pressure experiments, we used probit analysis of dose-response data collected at 30 minutes with the SPSS toolkit (IBM SPSS Statistics for Windows, Version 27.0). This analysis estimated the concentrations at which 25% and 75% of the AF populations were affected (LC_25_ and LC_75_) [21]. Based on these LC values, a subset of the AF population was subjected to insecticide pressure using insecticide-coated bottles as described previously [21,22]. Both males and females, aged 2-5 days, were included in these assays. For each insecticide (permethrin or lambda-cyhalothrin), about 950 mosquitoes were exposed to insecticide pressure in each filial generation (F1 or F2) at both LC_25_ and LC_75_ levels. Bioassays were performed in batches of 25 mosquitoes per bottle. To assess the impact of selective pressure, mortality rates were recorded for each filial generation at both LC_25_ and LC_75_.

### Larval bioassays for resistance validation

To assess changes in resistance levels in pyrethroid-selected strains, we conducted larval bioassays to obtain dose-response curves and estimate the LC_50_ and resistance ratios (RR) for lambda-cyhalothrin and permethrin following the WHO larval protocol [15]. Bioassays were conducted using late third or early fourth-instar larvae in jars containing 99 mL of water and 1 mL of insecticide. The control consisted of 1 mL of ethanol in 99 mL of water. Each experiment included two to three replicates, each with three repetitions, involving 20 larvae and five to seven increasing concentrations of insecticide. Permethrin concentrations ranged from 0.0015 to 0.0192 ppm, while lambda-cyhalothrin concentrations ranged from 0.001 to 0.1 ppm. Larval mortality was recorded 24 hours after exposure.

The dose-response data were used to calculate the lethal concentrations 50 and 90 (LC_50_ and LC_90_, respectively) for the unselected AF, and the second filial generation (F2) of the selected lambda-cyhalothrin-selected strains (HL and LL) and permethrin-selected strains (HP and LP). Resistance ratios 50 (RR_50_) and 90 (RR_90_) were calculated by dividing the LC_50_ and LC_90_ of the tested strains by those of the susceptible Rockefeller laboratory strain, respectively [23]. Significant differences between LC_50_ and LC_90_ were assessed using the lethal dose ratio test [21].

### Determination of insecticide resistance mechanisms in response to insecticide selection

After two generations of insecticide pressure, we examined potential resistance mechanisms between the unselected AF strain and the pyrethroid-selected strains (F2). We assessed changes in *kdr* allele frequencies, mean enzymatic activity through biochemical assays, and gene expression via transcriptomic analysis.

### *Kdr* allelic frequency

We employed an allele-specific PCR (AS-PCR) to detect the *kdr* mutations F1534C, V410L, and V1016I in the voltage-gated sodium channel gene. These mutations have been found in field populations of *Ae. aegypti* in Colombia and previously in our AF strain [24,25]. Genomic DNA was extracted from individual insects using the Grind Buffer protocol, which involves homogenizing each insect in 50 µL of buffer followed by incubation with 20 µL of proteinase K. The DNA was precipitated with potassium acetate, purified by centrifugation, and washed with 70% and 98% ethanol. The purified DNA was resuspended in 30 µL of water. PCR reactions were run on a Rotor-Gene Q thermocycler under conditions described by Pareja-Loaiza et al. (2020) [25]. The experiment included 30 adult mosquitoes from each of the pyrethroid-selected strains, the non-selected AF strain, and the Rockefeller reference strain as a control.

### Enzymatic bioassays

We examined metabolic resistance mechanisms by measuring enzyme activities in insecticide-selected mosquitoes from the F2 generation, the AF strain, and the Rockefeller reference population. The tested groups were labeled as HP, LP, HL, LL, and the AF population. Activities of acetylcholinesterase (AChE), alpha and beta esterases (α-EST and β-EST), mixed-function oxidases (MFO), and glutathione S-transferase (GST) enzymes were measured using established protocols [26]. Briefly, forty newly emerged female mosquitoes from each group were homogenized in 300 µL of deionized water with a tissue grinder and MicroPestle system. Enzyme activities were normalized to total protein concentrations, which were determined using the Pierce BCA Protein Assay Kit. The assays used specific substrates: DTNB for AChE (25 µL of homogenate with or without an inhibitor), TMBZ for MFO (20 µL), α-and β-naphthyl for α-EST and β-EST (10 µL of supernatant), and reduced glutathione for GST (15 µL of supernatant). Measurements were taken with the Thermo Fisher Scientific ELISA Multiskan Spectrum at previously reported wavelengths. The Kruskal–Wallis test was used to assess statistically significant differences among populations (p < 0.05). When appropriate, post hoc Dunn’s tests were performed to identify pairwise differences (p < 0.05) and enzyme activities were visualized using violin plots.

### Transcriptomic analysis

Transcriptomic analysis was performed using RNA sequencing (RNA-seq) to compare pooled individuals from the F2 pyrethroid-selected strains (Fig 1). To evaluate differences in gene expression (mRNA) between survivor and dead populations within the F2 offspring, we conducted an additional bioassay using the pyrethroid LC_75_ on those strains selected at high concentrations (HP and HL).

After a 30-minute exposure, survivors and dead mosquitoes were recorded as ‘resistant’ and ‘susceptible’, respectively. Mosquitoes from these pools were used to prepare libraries for lambda-cyhalothrin HL-susceptible (HLS), HL-resistant (HLR), and for permethrin-selected as HP-susceptible (HPS) and HP-resistant (HPR). In addition, we built libraries from F2 individuals of the LP and LL (LC_25_); however, these were not further split into survivor or dead populations due to the low number of ‘susceptible’ individuals (Fig 1). We examined transcriptomic changes 30 min after insecticide exposure, as previous evidence suggests that this duration is sufficient to induce transcriptomic modifications [27,28] and matches the exposure time originally used for pressure. RNA sequences were deposited in the Sequence Read Archive under the BioProject number PRJNA1273191.

RNA extraction was performed using the Spin Tissue RNA Mini Kit on three pools of five mosquitoes each from the selected HLR, HLS, LL, HPS, and LP populations and two pools from HPR. Mosquitoes were homogenized in lysis buffer using micropestles. RNA was eluted in 30 mL of buffer, and its quantity and integrity were checked by Nanodrop and electrophoresis gel, respectively. The RNA was then sent to Novogene Corporation Inc.

(Sacramento, CA) for sequencing. For comparison, we used RNA sequencing data from the unselected AF strain previously described [15].

We used FastQC [29] to assess the quality of RNAseq reads and Trimmomatic [30] to remove low-quality sequences with a Phred score below 30 and a length under 36 nucleotides. The AaegL5 genome, which contains 19,804 genes including 14,718 protein-coding genes, was used as a reference [31] for mapping using STAR [32]. Gene expression levels were quantified using the Rsubread package within DEseq2 [33]. Differentially expressed genes (DEGs) were identified by comparing the selected strains (HLR/HPR, HLS/HPS, LL, LP) with AF, using a log2 fold change (log2FC) of at least 2 and a False Discovery Rate (FDR) of < 0.05. The normalized data were then scaled for statistical analysis to facilitate group comparisons.

We reasoned that comparing resistant and susceptible mosquitoes within each LC would identify genes whose expression must be higher to survive the insecticide doses. Specifically, we focused on transcripts showing a stepwise increase in expression, upregulated at LC₂₅, further elevated in LC₇₅-susceptible mosquitoes, and peaking in LC₇₅-resistant individuals.

We identified DEGs with a log2FC expression pattern of HLR/HPR > HLS/HPS > LL/LP. This approach helped us focus on DEGs more likely associated with resistance and compare their expression across mosquito strains exposed to the same insecticide.

### Annotation with custom database and Gene Ontology (GO)

Gene annotations were performed using VectorBase. However, because of the large number of “unspecified products,” we identified additional annotations using a custom database, as Mack and Attardo noted [28]. Briefly, we extracted the protein FASTA files from NCBI txid 7214 (Drosophilidae) and txid 7157 (Culicidae). We created a merged database that added product descriptions to our VectorBase gene IDs. The query sequences of the “unspecified products” were replaced by the annotations provided by this custom database.

The R package TopGo was used for Gene Ontology enrichment analysis of our dataset [34]. A Fisher test with a p-value < 0.05 was used as the significance threshold. Figures were created using GraphPad version 8.0.0 for Windows (GraphPad Software, Boston, Massachusetts, USA, www.graphpad.com). Volcano plots were generated with R version 4.3.0 (December 2023), and upset plots were created using SRplot (https://www.bioinformatics.com.cn/en).

## Results

### The AF strain increased survival rates after insecticide pressure

First, we determined the diagnostic dose (DD) and diagnostic time (DT) for the AF strain of the insecticides lambda-cyhalothrin and permethrin by calculating mortality over time (Figs 2A and 2B). The DD and DT required to achieve 100% mortality were 24 ppm at 35 minutes for lambda-cyhalothrin and 150 ppm at 35 minutes for permethrin (Fig 2).

**Fig. 2.**
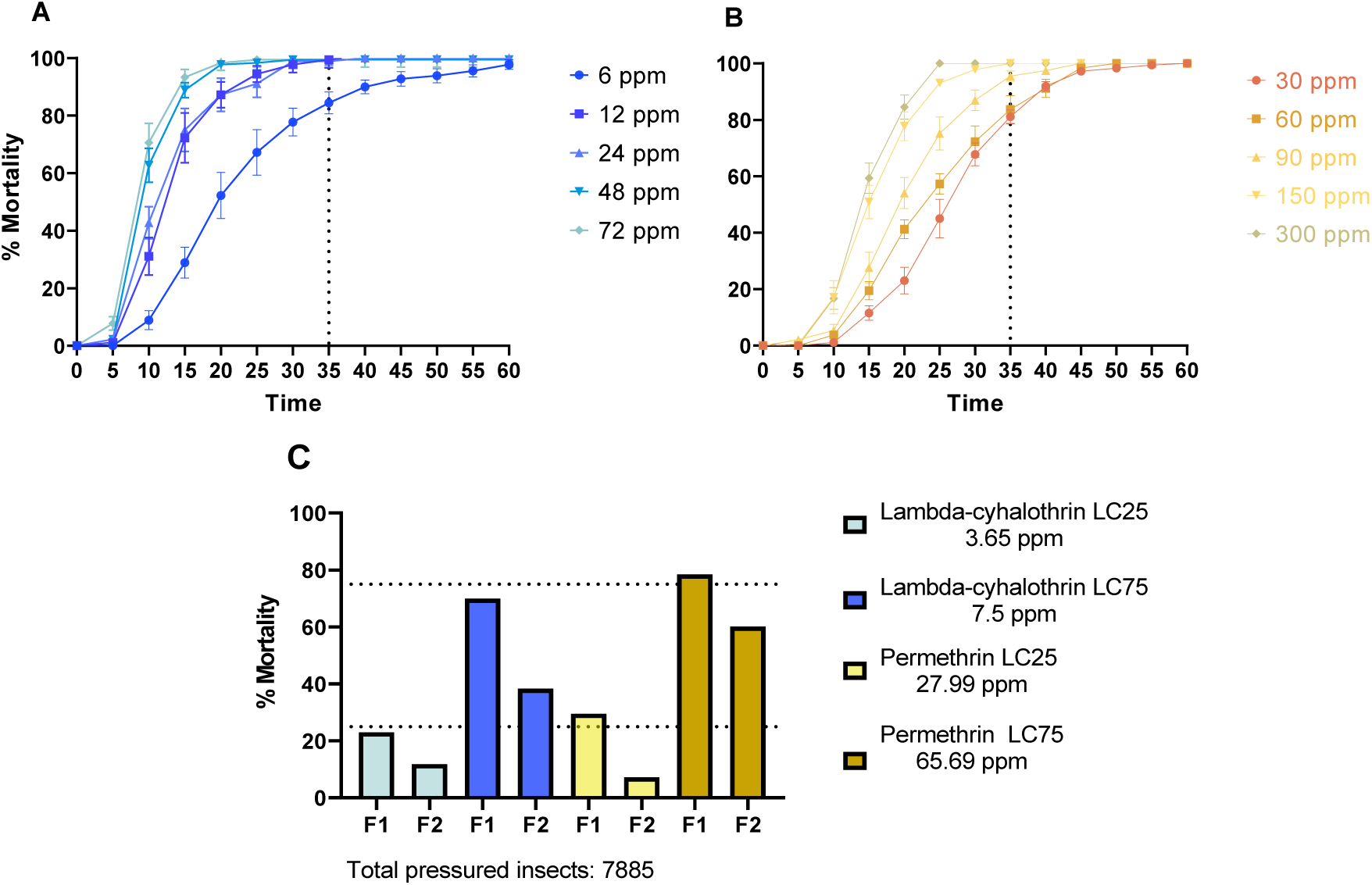
Response to insecticides in the AF population and pyrethroid-selected strains. The graph shows the percentage of mortality over time (minutes) for lambda-cyhalothrin (A) and permethrin (B). The vertical dashed line at each insecticide represents the 35-minute DT (C). Mortality of the F1 and F2 generations exposed to LC_25_ or LC_75_ for lambda-cyhalothrin (blue) and permethrin (yellow) was obtained from dose-response curves. The horizontal dashed lines correspond to the 25% and 75% mortality levels.

Then, we determined the lethal concentrations that killed 25% and 75% of mosquitoes (LC_25_ and LC_75_, respectively) after 30 minutes of exposure to selected resistant strains.

The LC_25_ values were 3.65 ppm for lambda-cyhalothrin and 27.99 ppm for permethrin; the LC_75_ values were 7.5 ppm for lambda-cyhalothrin and 65.69 ppm for permethrin (Fig 2C).

Subsequently, we exposed AF to these concentrations. As stated, the survivors of the LC_25_ were labeled low-pressure (L) strains, and those of the LC_75_ were labeled high-pressure (H) strains. Changes in susceptibility were evaluated by comparing the mortality responses to these concentrations in offspring from the first (F1) and second (F2) selections (Fig 2C).

In the selected F1 generation, lambda-cyhalothrin mortality was 23% and 70% for the strains selected at LC_25_ (LL) and LC_75_ (HL), respectively. For permethrin, mortality was 29.5% and 78.4% for the strains selected at LC_25_ (LP) and LC_75_ (HP), respectively (Fig. 2C).

In the F2 selected generation, lambda-cyhalothrin mortality was 11.8% and 38.3% for LL and HL, respectively. For permethrin, mortality was 7.2% and 60.1% for LP and HP, respectively. Overall, we observed a significant increase in mosquito survival after two generations of selection (compared with the AF population). Therefore, we used the F2 generation to evaluate potential mechanisms of insecticide resistance.

### Pyrethroid-selected strains have higher resistance ratios than the AF strain

We applied pyrethroid selective pressure on adult mosquitoes of both sexes using the bottle bioassay. However, to validate changes in resistance, we performed a larval bioassay following WHO guidelines, obtaining LC_50_ and LC_90_ values for each strain (Fig 1 and Table 1). These values were used for relative comparisons by calculating resistance ratios (RR_50_ or RR_90_) between the lambda-cyhalothrin-selected F2 strains (HL and LL), permethrin-selected F2 strains (HP and LP), the unselected AF, and the susceptible reference Rockefeller strain (Table 1).

**Table 1.** Insecticide pressure induces changes in the resistance ratios (RR) of selected *Ae. aegypti* strains. Resistance ratios to lambda–cyhalothrin in larva of Acacías and Rockefeller strains.

| *Ae. aegypti* strain | Generation | N | Pressure | LC<sub>50</sub> (ppm) | (95% CI) | LC<sub>90</sub> (ppm) | (95% CI) | RR<sub>50</sub> | RR<sub>90</sub> | X<sup>2</sup> | df | Slope ± SE |
| --- | --- | --- | --- | --- | --- | --- | --- | --- | --- | --- | --- | --- |
| Rockefeller | N/A | 1260 | N/A | 0.00047 | (0.000398–0.000564) | 0.0016 | (0.0012–0.0022) | 1.0 | 1.0 | 65.0 | 61 | 2.0 ± 0.1 |
| AF | 0 | 840 | No | 0.005 | (0.003–0.007) | 0.017 | (0.014–0.21) | 10.5** | 10.6** | 21.0 | 40 | 2.4 ± 0.44 |
| LL | F<sub>2</sub> | 620 | Low (LC<sub>25</sub>) | 0.007 | (0.004–0.011) | 0.044 | (0.023–0.424) | 14.8** | 27.5** | 296.0 | 30 | 2.5 ± 0.21 |
| HL | F<sub>2</sub> | 600 | High (LC<sub>75</sub>) | 0.061 | (0.031–0.168) | 0.157 | (0.1–0.659) | 129.7** | 98.1** | 333.2 | 28 | 13.3 ± 1.5 |

Resistance ratios to permethrin in larva of Acacias and Rockefeller strains.
| *Ae. aegypti*<br>strain | Generation | N | Pressure | LC<sub>50</sub><br>(ppm) | (95% CI) | LC<sub>90</sub><br>(ppm) | (95% CI) | RR<sub>50</sub> | RR<sub>90</sub> | X<sup>2</sup> | df | Slope<br>± SE |
| --- | --- | --- | --- | --- | --- | --- | --- | --- | --- | --- | --- | --- |
| Rockefeller | N/A | 1260 | N/A | 0.00022 | (0.000192–<br>0.000244) | 0.0008 | (0.000670–<br>0.000821) | 1 | 1 | 212 | 61 | 2 ±<br>0.1 |
| AF | 0 | 1260 | No | 0.028 | (0.023 –<br>0.035) | 0.065 | (0.055– 0.081) | 127.3** | 81.7** | 298.5 | 61 | 34.6<br>± 1.9 |
| LP | F<sub>2</sub> | 840 | Low<br>(LC<sub>25</sub>) | 0.038 | (0.03 – 0.049) | 0.078 | (0.063– 0.106) | 172.7** | 98** | 244.2 | 40 | 31.8<br>± 2.0 |
| HP | F<sub>2</sub> | 720 | High<br>(LC<sub>75</sub>) | 0.082 | (0.069–0.103) | 0.144 | (0.119–0.190) | 372.7** | 180.9** | 111.9 | 34 | 20.6<br>± 1.7 |

Bioassays were used to determine the LC<sub>50</sub> and LC<sub>90</sub>, with 95% confidence intervals (CIs), for each strain. The (RR) was obtained by dividing the LC of the target strain by that of the Rockefeller strain. \*\*Resistant strains, N, number of larvae used in the bioassay; N/A, not applicable; SE, standard error. The LC<sub>50</sub> and LC<sub>90</sub> values for all pressured strains were significantly different compared to the Rockefeller strain ( $p < 0.05$ ).

These values were used for relative comparisons by calculating resistance ratios (RR50 or RR90) among the lambda-cyhalothrin-selected F2 strains (HL and LL), permethrin-selected F2 strains (HP and LP), the unselected AF, and the susceptible reference Rockefeller strain. The results show that both the AF strain and those treated with both insecticides at the two concentrations are resistant (Table 1).

Comparing the pyrethroid-selected strains with the unselected AF allows us to assess the response to artificial selection. For lambda-cyhalothrin, RR_50_ increased 12.3-fold in HL and 4.3-fold in LL. For permethrin, RR_50_ increased 2.9-fold in HL and 1.4-fold in LL. Although the 95% confidence intervals for the low-concentration-selected strains overlap with AF, significant differences were observed relative to the Rockefeller strain (p < 0.05). Additionally, significant differences (p < 0.05) were observed between the AF strain and the high-concentration-selected strains (HP and LP).

Overall, the selection pressure from AF resulted in four strains with distinct insecticide susceptibility profiles, which were then used to investigate the mechanisms of insecticide resistance.

### Allelic frequencies showed minor changes in response to two rounds of insecticide pressure

We evaluated the effect of pyrethroid selection on the frequency of *kdr* alleles. We tracked changes in genotype frequencies at three loci: V410L, V1016I, and F1534C, in our strains (Fig 1). For this analysis, we used HP and HL mosquitoes exposed to LC_75_ and LP, and LL mosquitoes exposed to LC25 (Fig 3).

**Fig. 3.**
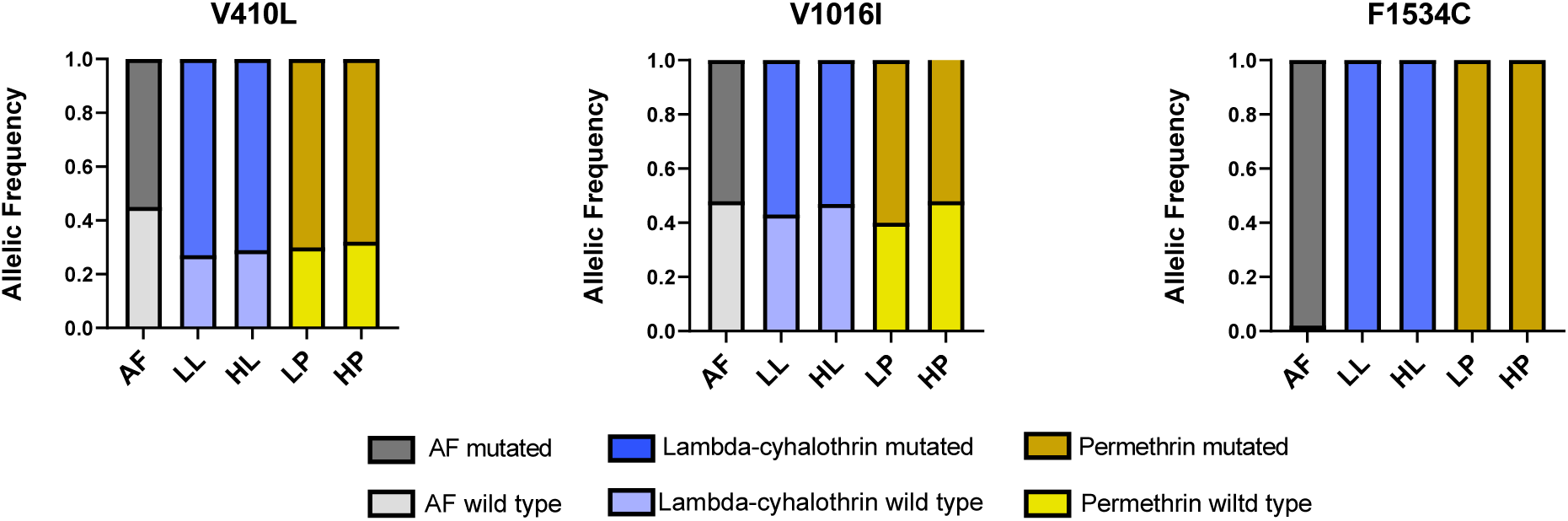
*kdr* allelic frequencies in strains exposed to lambda-cyhalothrin and permethrin, as well as the unselected AF. The dark color corresponds to the mutated alleles 410L, 1016I, and 1534C. The light color corresponds to the wild-type alleles V410, V1016, and F1534. Blue and yellow depict lambda-cyhalothrin and permethrin, respectively. The unpressured AF population is shown in gray. Dark gray represents the mutated allele, while light gray represents the wild-type allele.

The V410L mutant allele had frequencies of 0.68 to 0.73 in the pressured strains, representing an increase of 0.13 to 0.18 over the 0.55 observed in the AF strain. Specifically, permethrin-pressured strains had frequencies of 0.7 and 0.68 for LP and HP, respectively, while lambda-cyhalothrin-pressured strains had frequencies of 0.73 and 0.71 for LL and HL, respectively.

The frequencies of the 1016I allele were 0.53 in both the high lambda-cyhalothrin-and permethrin-pressure strains. In the low lambda-cyhalothrin-and permethrin-pressure strains, the frequencies were 0.57 and 0.6, respectively. The control AF strain showed an allele frequency of 0.52. This represented a non-significant increase in mutated allele frequency, ranging from 0.1 to 0.7.

Analysis of F1534C allele frequencies showed that the 1534C mutation was dominant in treated strains, with a wild-type allele frequency of only 0.02 in the AF strain. This indicates that after 2 generations of insecticide pressure, even at a low concentration, the mutated allele becomes fixed in the population.

Overall, we observed only subtle differences in *kdr* allele frequencies among our strains in response to varying concentrations of permethrin and lambda-cyhalothrin. Thus, the mutated alleles in these insecticide-pressure strains are unlikely to account for the previously observed differences in RR (Table 1).

### Enzyme activity is altered in mosquitoes exposed to permethrin

One of the primary mechanisms by which pyrethroid insecticides are detoxified is enzyme-mediated degradation. The biochemical assays in our study reflect broad enzyme activity towards model substrates and are not specific to the metabolism of insecticides. Nevertheless, they provide a useful indication of potential metabolic resistance. Enzymatic activity assays were performed on the F2 generations of LP, HP, LL, and HL (Fig. 4). To evaluate differences in activity caused by pyrethroid selection, we compared the high-and low-selection strains with the unselected AF strain.

**Fig. 4.**
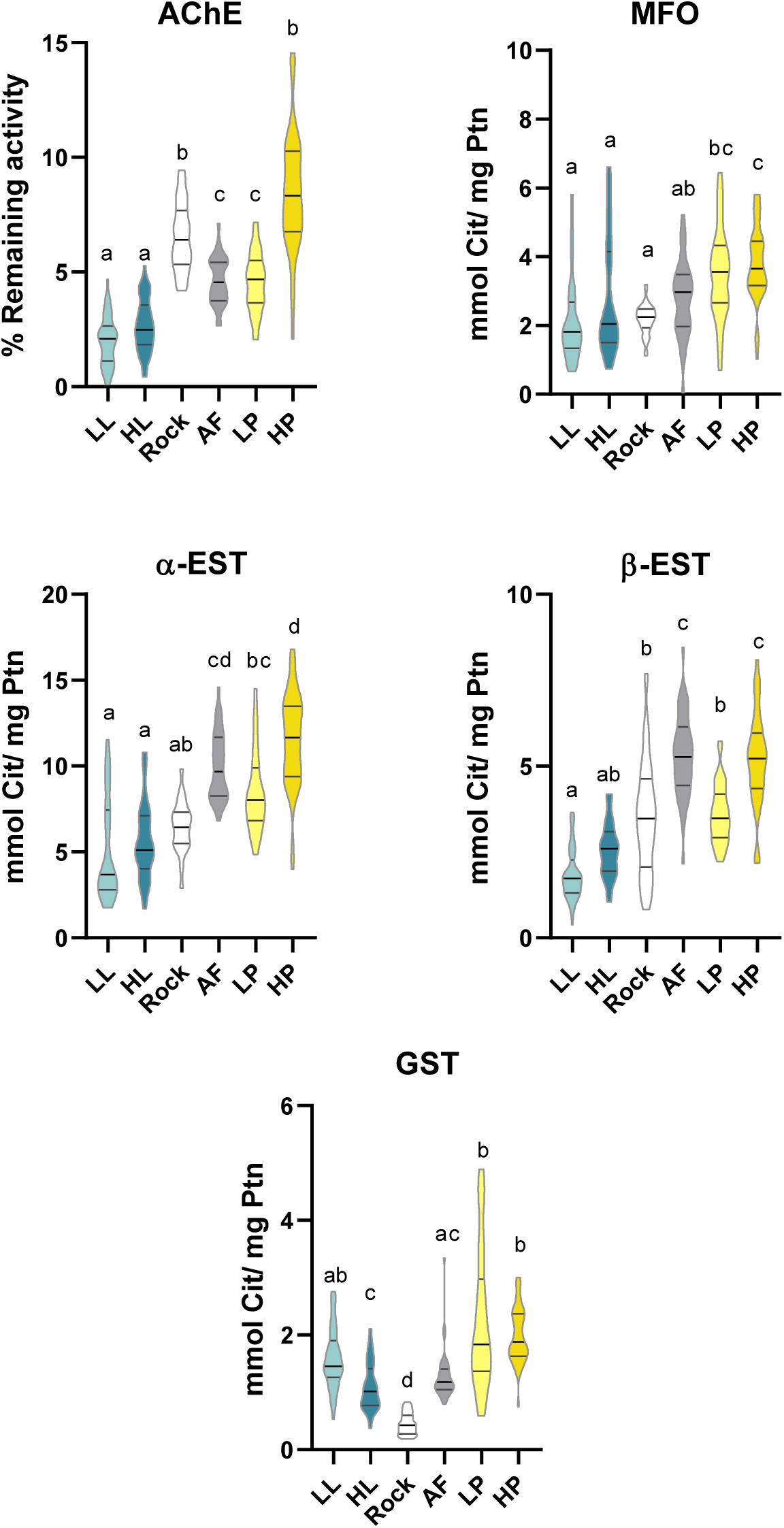
Enzymatic activity in pyrethroid-selected strains (HL and LL selected with lambda-cyhalothrin; HP and LP selected with permethrin), the unselected strain (AF), and the susceptible reference strain Rockefeller. Enzyme activity was measured using model substrates and categorized as AChE (acetylcholinesterase), MFO (mixed-function oxidase), α-EST (alpha-esterases), β-EST (beta-esterases), and GST (glutathione S-transferase). Violin plots represent the distribution of individual values; central lines indicate the median, and boxplots show the interquartile range. Significant differences among strains were determined using Kruskal–Wallis tests followed by Dunn’s multiple comparison test with Bonferroni correction. Specific pairwise comparisons are provided in S1 Table.

No significant differences were detected between the lambda-cyhalothrin-selected strains (HL and LL) for AChE, MFO, α-EST, or β-EST activities (p > 0.05). However, GST activity was higher in LL than in HL (Fig. 4). However, HL and LL strains had significantly higher GST activity than Rockefeller, whereas mean AChE, α-EST, and β-EST activities were significantly lower in both HL and LL strains than in AF.

In the permethrin-selected HP and LP strains, AChE, α-EST, and β-EST activities were significantly higher in HP than in LP. When HP was compared with the unselected AF, higher AChE, MFO, and GST activities were observed in HP. Comparisons between LP and AF showed that the mean GST activity was higher in LP, while mean β-EST activity was lower.

Overall, the response of the lambda-cyhalothrin-selected strains differed from that of the permethrin-selected strain. In lambda-cyhalothrin-selected populations, significant differences were observed in AChE, α-EST and β-EST, which tended to be lower compared with the reference or unselected strains. Conversely, in permethrin-selected populations, AChE, α-EST and β-EST activities tended to be higher in HP than in LP, as did MFO and GST activities when compared with AF. These results suggest differential enzymatic response patterns associated with insecticide concentration. However, there is insufficient data to conclude that the complex interactions among these phenotypes and insecticide resistance can be explained solely by differences in enzymatic activity.

### Transcriptomic analyses

As no significant differences were observed among the common resistance mechanisms (e.g. *kdr* allele frequencies or enzymatic activity), our focus was on identifying differences in gene expression in our pyrethroid-selected strains. At this stage, we included the susceptible individuals that did not survive the LC₇₅ exposure. These individuals are referred to as HPS and HLS for permethrin and lambda-cyhalothrin, respectively. This approach allowed us to concentrate on transcripts showing a stepwise increase— rising from LC₂₅ to LC₇₅-susceptible and peaking in LC₇₅-resistant mosquitoes — consistent with transcriptional responses associated with pyrethroid resistance.

To explore these expression patterns, we performed RNA-seq analysis. After sequencing and trimming, we obtained 758,574,050 reads, which we aligned to the *Ae. aegypti* genome (version LVP_AGWG AaegL5.3) using STAR. This produced approximately 91.6% unique reads. Of these, 70.5% mapped uniquely, and 20.2% mapped to multiple loci (S2 Table). We used a log2FC threshold of 2 to identify differentially expressed genes (DEGs) (Figs 5-8).

**Fig. 5.**
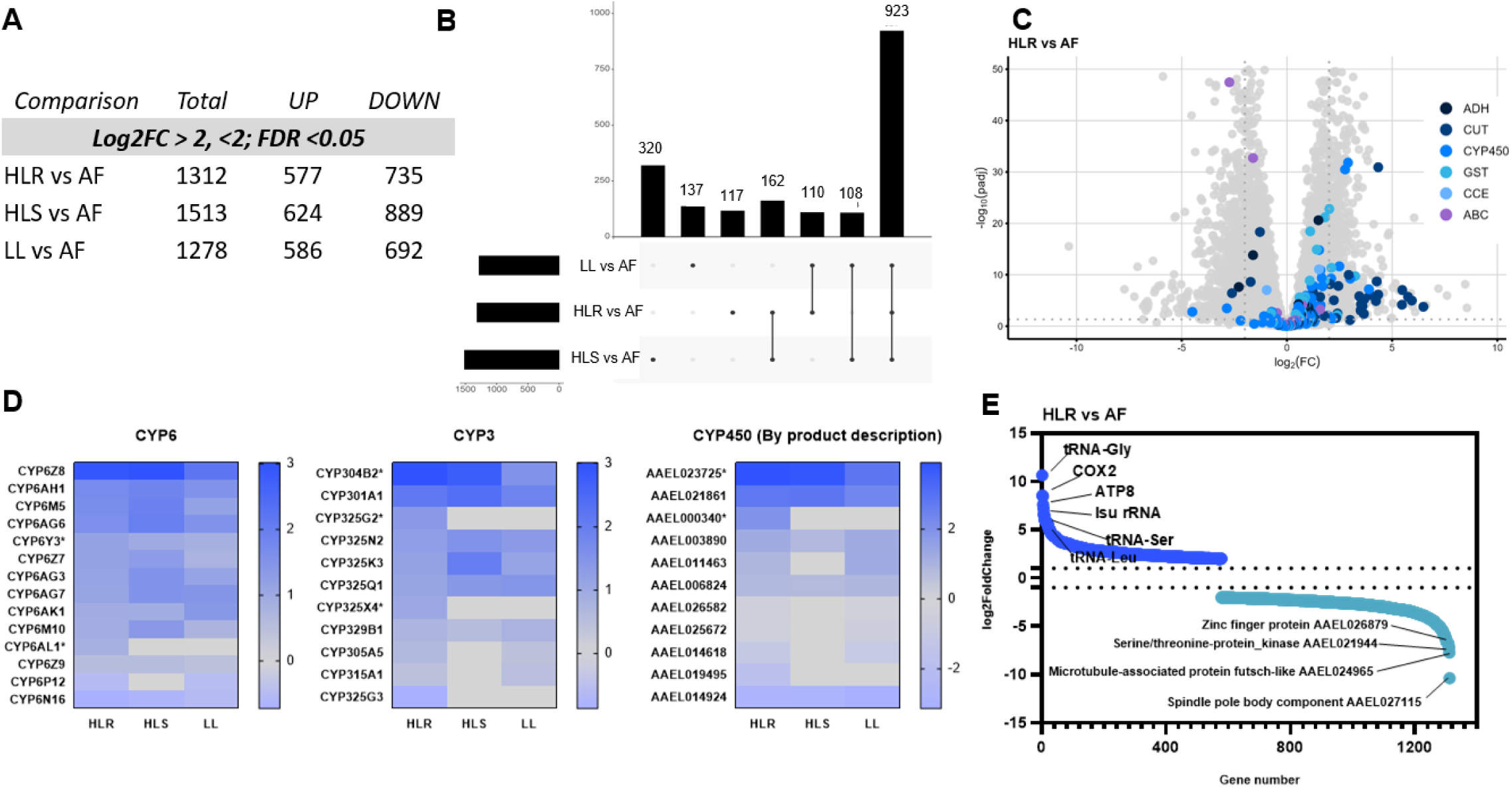
Transcriptomic analysis stratified by IR genes and the top-regulated DEGs in lambda-cyhalothrin-treated strains HLR, HLS, LL, and their comparison with AF. (A) The number of DEGs up-or down-regulated across the strains. (B) Upset plot of unique and shared genes among the pressured strains and AF. (C) Volcano plot of the main comparison HLR vs AF; ADH, aldehyde deoxygenase; CUT, cuticle; CYP450, cytochrome P450; GST, glutathione S-transferase; CCE, carboxy/choline esterase; ABC, ABC transporters. (D) Gene categories of the main CYP subfamilies whose log2 Fold Change (Log2FC) were HLR > HLS > LL or HLR > HLS/LL. (*) Represents the genes that met this condition. The log2FC is depicted by the blue gradient. (E) Cascade plot of the top up-and downregulated genes in the HLR vs AF comparison.

**Fig 6.**
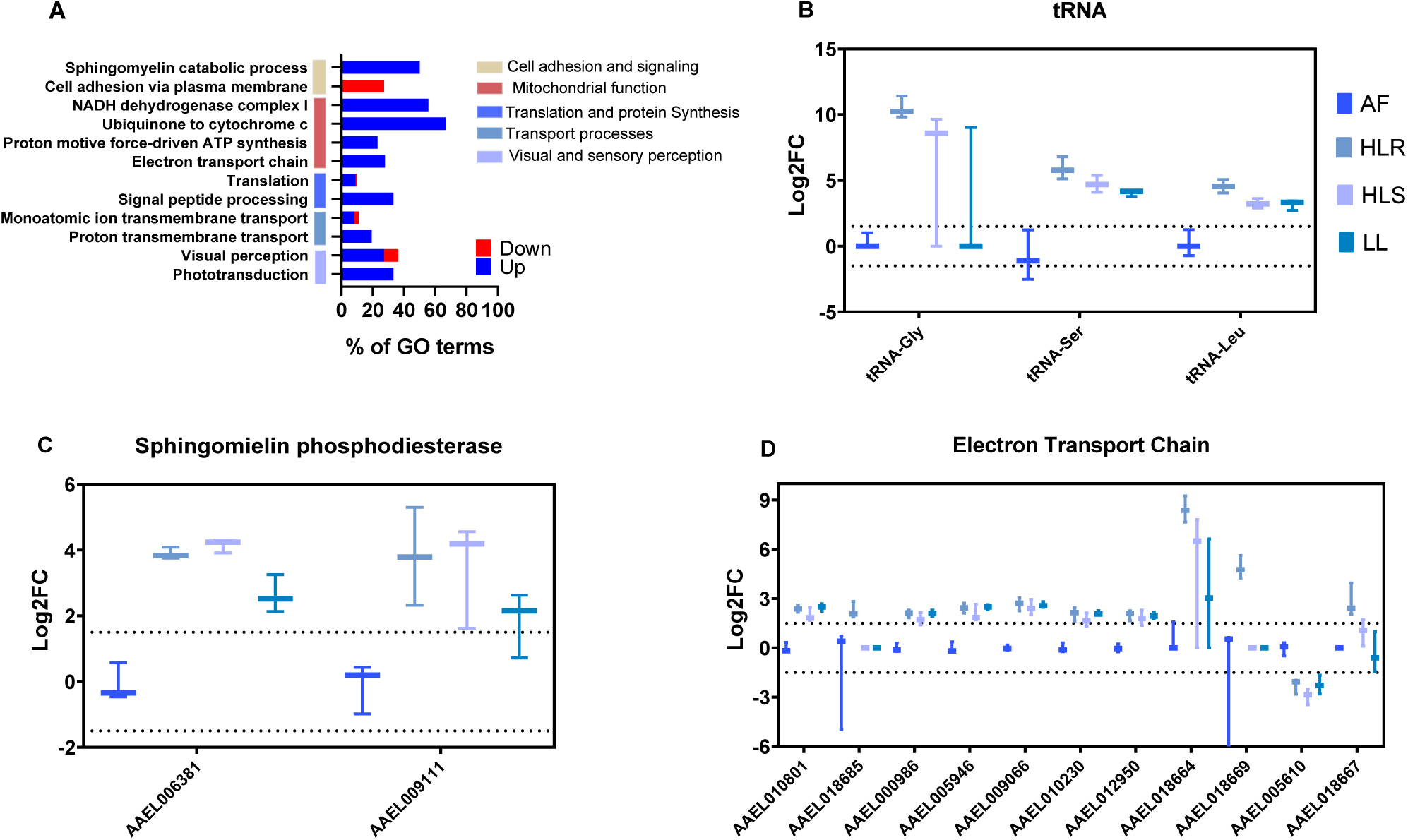
Lambda-cyhalothrin pressure induces a consistent upregulation of translation processes and cellular respiration. (A) Gene term enrichment analysis of DEGs (HLR vs AF) shows the percentage of upregulated (blue) and downregulated (red) genes in our data. The length represents the proportion of regulated genes within each term, expressed as a percentage. Vertical colored lines indicate gene classifications, with their corresponding legends. (B) Bar plot of the most upregulated tRNA genes that are present in Fig 5E. (C and D) Bar plots of significant genes involved in cell adhesion and mitochondrial function, respectively. Horizontal lines represent log2FC values greater than 2. Only genes with an FDR < 0.05 are shown in the bar plots.

**Fig 7.**
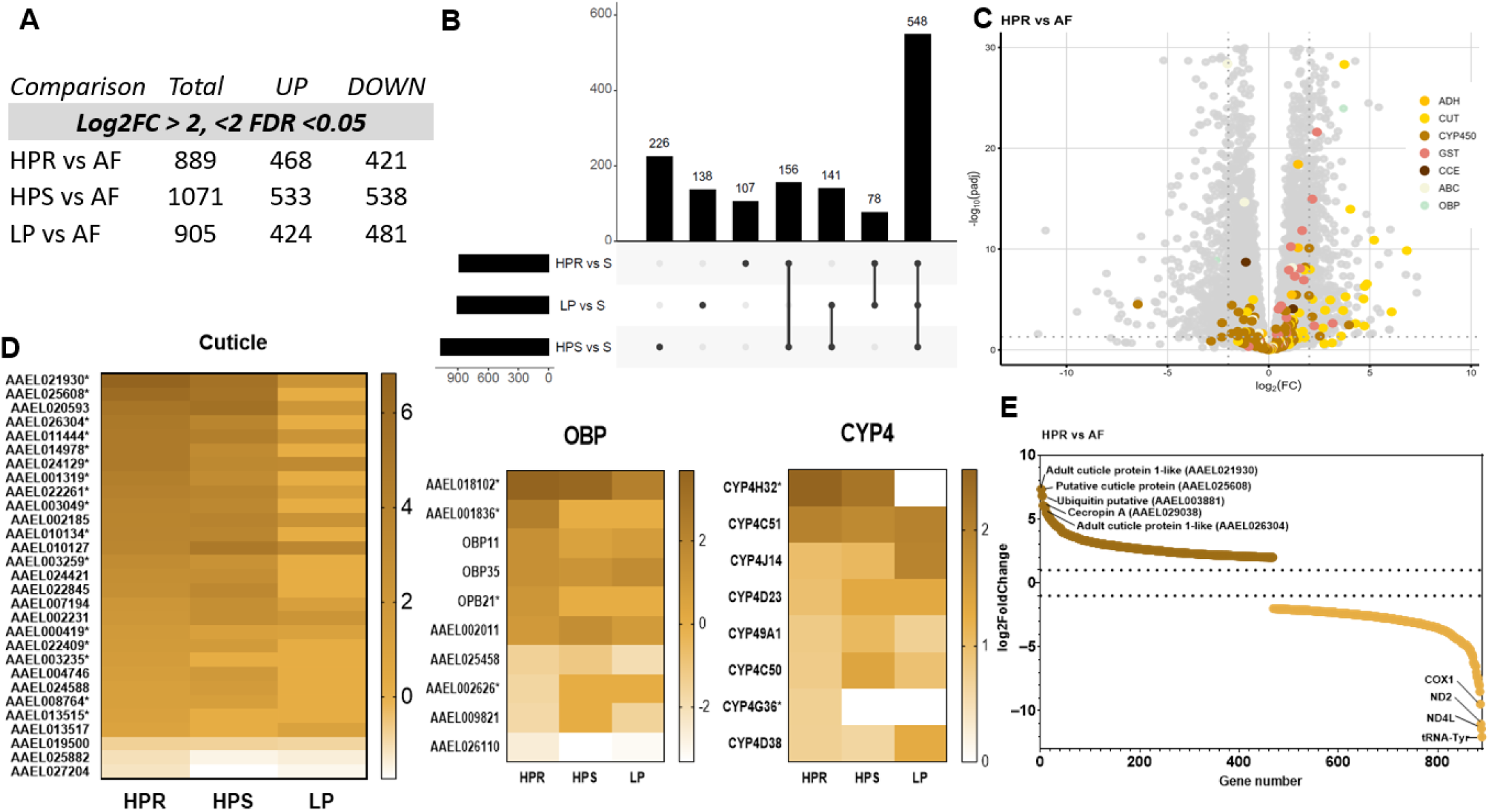
Transcriptomic analysis of representative insecticide resistance-related genes in permethrin-treated populations and AF, showing the overregulation of most up-and down-regulated genes presented in this data set. (A) Number of differentially expressed genes (DEGs) that are up-or down-regulated among the populations compared with AF. (B) Upset plot of the unique and shared genes between pressured strains and AF. (C) Volcano plot of the comparison HPR vs AF; ADH, aldehyde deoxygenase; CUT, cuticle; CYP450; cytochrome P450; GST, glutathione S-transferase; CCE, carboxy/choline esterase; ABC, ABC transporters and OBP, Odorant Binding Proteins. (D) Gene categories of the main cuticle, OBP, CYP4 whose log2 Fold Change (Log2FC) was HPR > HPS > LP or HPR > HPS/L. (*) Represents the genes that follow this condition. Log2FC is depicted by the yellow gradient color. (E) Cascade plot of the top-up and downregulated genes in the HLR vs AF comparison.

**Fig 8.**
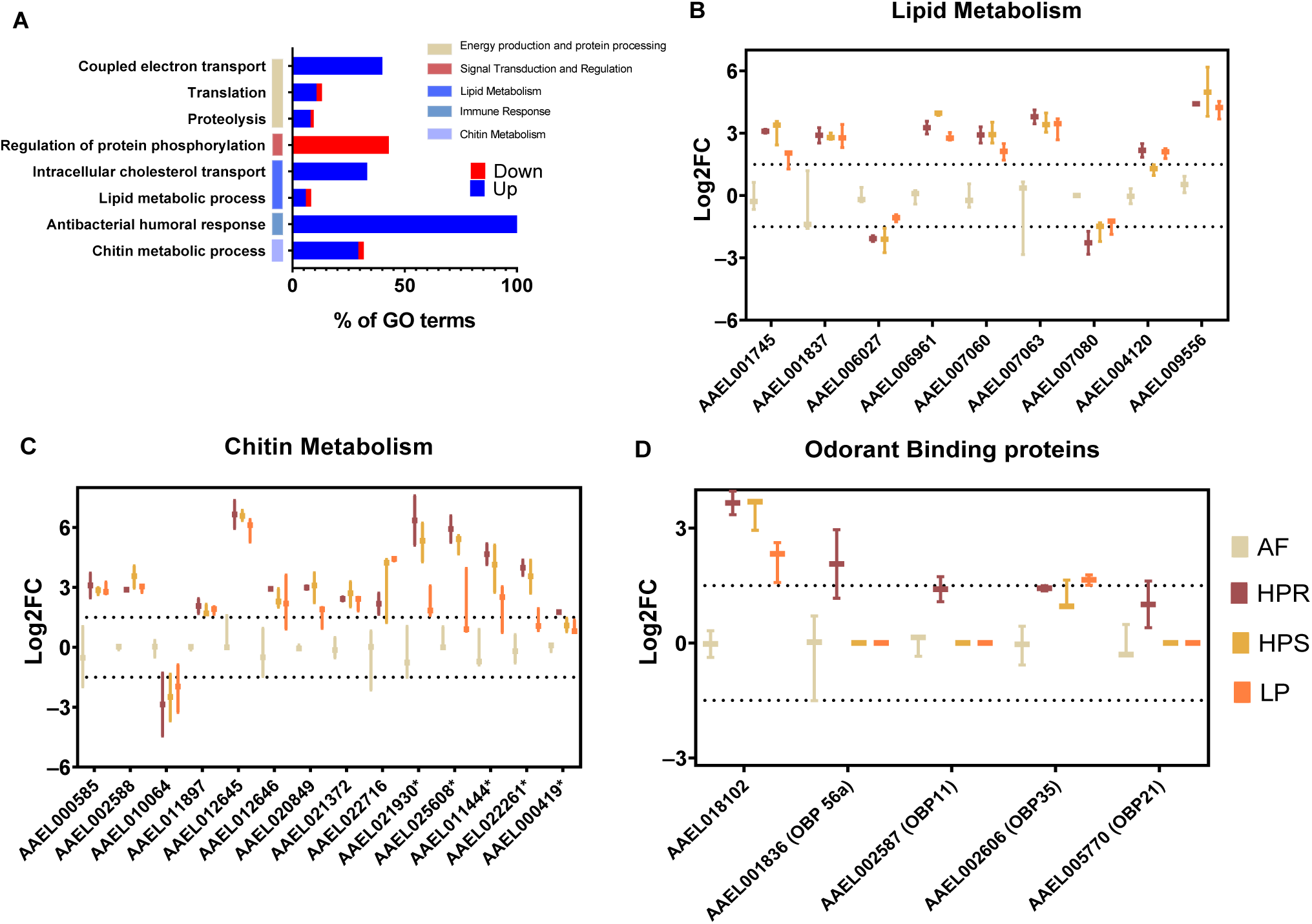
Percentage of GO terms related to permethrin exposure at high concentrations and individual gene log2FC expression in categories associated with GO terms. (A) Gene term enrichment analysis of DEGs (HPR vs AF), showing the percentage of upregulated (blue) and downregulated (red) genes in our data. The length represents the proportion of regulated genes within each term, expressed as a percentage. Vertical colored lines indicate the classification of genes, accompanied by their corresponding legends. Bar plot of genes related to (B) lipid metabolism (C), chitin metabolism, and (D) odorant binding proteins. Horizontal lines represent log2FC values greater than 2. Only genes with an FDR < 0.05 are showcased in the bar plots.

### tRNA, electron transport chain, CYP450 leads the pattern of upregulation related to lambda-cyhalothrin pressure

We analyzed the transcriptome of insects exposed to lambda-cyhalothrin to understand its relationship to resistance at different concentrations of the insecticide. We compared the transcriptomes of the HLR, HLS, and LL populations with the AF population to identify unique and shared DEGs (Figs 5A and 5B). A total of 4,103 DEGs were identified, 923 of which were present in all the selected populations. 320 DEGs were exclusive to the HLS population, 137 to the LL population, and 117 to the HLR (Fig 5B). Our analysis focused on 203 differentially expressed IR-related genes, including cytochrome P450, cuticle, glutathione S-transferase, carboxylesterase/choline esterase, and ABC transporters.

Among these 203 DEGs, we searched for those common across the selected populations and whose expression followed the pattern (HLR > HLS > LL). Only 17 (8.3%) of the 203 DEGs matched this pattern, with 13 belonging to the CYP450 family and the remaining four being GST and ABC transporters. The remaining 186 DEGs (91.7%) were related to IR but did not follow the pattern (HLR > HLS > LL) (Fig 5D). However, they were all upregulated in HLR, HLS, and LL compared to AF. The majority of these (145/203 = 71%) were related to cytochrome P450 and cuticle-related genes (Fig 5C and 5D).

Accordingly, we examined genes with the highest levels of up-and downregulation in the HLR population (Fig 5E). The top upregulated DEGs included three tRNAs (tRNA-Gly, tRNA-Ser, and tRNA-Leu), two electron transport genes (COX8 and ATP8), and one ribosomal RNA subunit (LSU rRNA), with log2FC ranging from 6.04 to 10.6. This corresponds to a Fold Change of 60 to 1000 times, respectively, compared with control. The most downregulated genes included a spindle pole body component (AAEL027115), a microtubule-associated protein (AAEL024965), a serine/threonine kinase (AAEL021944), and a zinc finger protein (AAEL026879), with log2FC between 6.7 and 10.3 (Fig 5E and S3 Table).

Some interesting genes that did not surpass the stringent threshold (log2FC > 2) are listed in S3 Table. These include several CLIP domain serine proteases from families A, B, C, and D, as well as four caspases (CASPS17, Dredd, CASPS8, and CASPS19) with log2FC > 1.

We then performed gene ontology analysis on the up-and down-regulated DEGs (1,312 genes) in HLR vs AF and compared them with the 11,450 genes from the entire dataset (Fig 6A). We identified five distinct biological processes, including cell adhesion and signaling, mitochondrial function, translation and protein synthesis, transport and vision, and sensory perception (Fig 6A). Among these, 12 terms were shared across these processes: phototransduction, visual perception, proton transmembrane transport, monoatomic ion transmembrane transport, signal peptide processing, translation, electron transport chain, proton motive force-driven ATP synthesis, ubiquinone to cytochrome C, NADH dehydrogenase complex I, cell adhesion via the plasma membrane, and sphingomyelin catabolic process. Most of these terms were upregulated in HLR (Fig 6), except for cell adhesion via the plasma membrane.

We compared the log2FC of selected genes across the HLR, HLS, LL, and AF strains, focusing on tRNA, sphingomyelin phosphodiesterase, and the electron transport chain (Fig 6B, C, and D). These genes showed the greatest upregulation and were also identified in the GO analysis.

Regarding mitochondrial function, we observed 24 upregulated genes. Specifically, those with a log2FC above 2 included AAEL010801 (Cytochrome b-c1), AAEL018685 (CYTB), AAEL000986 (NADH-ubiquinone oxidoreductase ashi subunit), AAEL005946 (NADH-ubiquinone oxidoreductase subunit B14.5b), AAEL009066, AAEL010230 (NADH ubiquinone dehydrogenase), and AAEL012950 (NADH-ubiquinone oxidoreductase). Interestingly, AAEL018664 (COX2), AAEL018669 (COX3), AAEL005610 (mitochondrial ATP synthase b chain), and AAEL018667 (ATP8) followed our pattern (HLR > HLS > LL) (Fig 6D).

Another term that is overrepresented in this dataset relates to translation and signal peptide processing. Several of these genes show the greatest upregulation (Fig 5E) and are associated with these processes (Fig 6A), including tRNAs. The expression of these three tRNAs followed the pattern (HLR > HLS > LL) (Fig 6B). Two other genes associated with sphingomyelin (AAEL006381 and AAEL00911) were upregulated in the HLR population and were emphasized due to their link to the nervous system and GO analysis.

### CYP, OBP, and cuticle genes are upregulated in insects resistant to high concentrations of permethrin

The transcriptome of permethrin-exposed populations revealed 2,865 differentially expressed genes (DEGs) across all treated groups compared with AF (Fig 7A). 548 genes were present in all samples (HPR, HPS, and LP) (Fig 7B). Additionally, the search for IR-genes in the HPR vs AF comparison identified 250 related to this phenotype. Among them, 156 were associated with cytochrome P450 and cuticle, 38 with OBP, 27 with GST, 18 with ABC transporters, 7 with ADH, and 4 with CCE (Fig 7C). This transcriptome showed clear upregulation of these genes, with 143 upregulated and 81 showing p-adjusted values above the threshold, whereas 107 were downregulated and only 31 exceeded the p-adjusted threshold (Fig 7C).

We found that 27 genes (10.8%) displayed the pattern (HPR > HPS > LP), and 16 of these (6.4%) were related to the cuticle. Most genes in these comparisons (89%) did not follow this pattern.

By analyzing the most up-and downregulated DEGs, we identified genes involved in cuticle formation and energy metabolism (Fig. 7E). The adult cuticle genes AAEL021930 and AAEL025608 were the two most upregulated in the dataset, with log2FC values of 6.8 and 6.07, respectively. They were followed by the ubiquitin gene (AAEL003881) and cecropin A (AAEL0290038), with log2FC values of 6 and 5.9, respectively. Another highly expressed cuticle gene, AAEL020593, had a log2FC of 5.2.

Genes that were most downregulated included a tRNA-tyr and three electron transport chain genes: ND4L, ND2, and COX1, with log2FC values of 12, 11.4, 9.4, and 8, respectively (Fig 7E).

Enzymatic bioassays revealed increased MFO activity in insects exposed to permethrin (Fig. 3). We identified genes from this family (CYP450); however, they do not account for differences in resistance and susceptibility at high or low concentrations. We conducted GO analysis to identify overrepresented gene terms in HPR that might explain these differences.

For the GO analysis, we assessed whether genes were predominantly upregulated or downregulated across each GO biological process (Fig 8A). The biological processes enriched among upregulated genes were grouped into four categories: energy production and protein processing, lipid metabolism, immune response, and chitin metabolism. The downregulated terms were related to signal transduction and protein phosphorylation regulation.

We examined the expression of each DEG in GO terms related to lipid and chitin metabolism, which are well-represented in the overall transcriptome and in the OBP genes (Figs 7B, 7C, and 8D). Regarding lipid metabolism, we found no clear differences among HPR, HPS, and LP, only an increase compared to AF. In contrast, some chitin metabolism genes followed a pattern (HPR > HPS > LP). These included AAEL021930, AAEL025608, AAEL011444, and AAEL022261 (Fig 8C). The OBP DEGs were upregulated in HPR, especially AAEL018102, OBP56a, OBP11, and OBP21.

In summary, lambda-cyhalothrin pressure led to the upregulation of genes involved in mitochondrial function and protein synthesis, whereas permethrin mainly induced transcripts related to immune response, lipid and chitin metabolism, in addition to the typical IR-related products. Although genes involved in the electron transport chain were affected by both insecticides, they were upregulated in lambda-cyhalothrin-pressured insects and downregulated in permethrin-pressured insects. The expression pattern we initially established was observed in lambda-cyhalothrin-pressured insects, characterized by the upregulation of transcripts for tRNAs and CYP450 enzymes. However, a different pattern was observed with permethrin, in which lipid metabolism was unaffected by this pattern, while other genes related to OBP and cuticle followed insecticide concentration. Overall, these findings support the observation that certain transcripts may be required at higher levels to resist the impact of increasing insecticide doses, and that success – or failure – to do so could be the difference between “resistance” and “susceptibility” in this strain.

## Discussion

In this study, we investigated the resistance mechanisms of *Aedes aegypti* following exposure to two concentrations of the pyrethroid insecticides lambda-cyhalothrin and permethrin over two generations. The mechanisms of insecticide resistance (IR) are complex and often intertwined. However, under selection pressure, the number of resistant insects increases while the number of susceptible insects declines [35], with distribution patterns depending on insecticide concentrations [12]. We tracked *kdr* mutations, conducted enzymatic activity assays to test for metabolic resistance, and performed transcriptomics to generate a dataset for comparing pressured and non-pressured mosquitoes. We employed an incremental overregulation approach (High Resistant > High Susceptible > Low Resistant) to identify differentially expressed genes (DEGs). These biological features are discussed in the following sections.

### Some *Kdr* allele frequencies are unaffected by insecticide pressure

We observed an opposing pattern, whereby insecticide pressure does not affect the frequency of mutant alleles, which remain consistent with those in the AF population. This could reflect the fact that fitness costs associated with *kdr* alleles may vary by allele and population [36].

Nonetheless, the presence of these alleles does not always correlate with IR in some populations [37]. One reason for this is that maintaining resistance-conferring alleles, such as 410L+1016I+1534C, is biologically more costly than maintaining only 1534C in the presence of an insecticide [38]. Thus, our data suggest that strains exposed to higher concentrations have lower frequencies of mutant alleles, possibly because the biological cost reduces the fitness of, or leads to the death of, triple-mutant individuals. Further investigation is warranted to confirm this hypothesis, as mutations alone may not be able to overcome high insecticide concentrations. It is assumed that the 410L mutation provides physiological benefits when paired with the 1534C mutation; however, this mutation is present in both resistant and susceptible insects [39]. In summary, we believe that in these strains, 1534C and 410L act as a “buffer” to maintain stable IR. However, under pressure, other mechanisms may develop, particularly in survivors of high insecticide concentrations.

### Permethrin high pressure but not lambda-cyhalothrin, enhances MFO activity

Following exposure to a xenobiotic, one of the initial responses in an insect is an increase in detoxification processes [40]. MFO is a well-known group of enzymes that convert pyrethroids into excretable metabolic intermediates [7]. In this study, we demonstrate a significant increase in MFO activity in the permethrin-selected HP strain relative to the AF and Rockefeller strains, and a slight increase in the LP strain relative to the AF strain. However, despite having a high resistance ratio (RR) to lambda-cyhalothrin, MFO levels in the HL and LL pressured strains are low. There are reports of resistance to type I insecticides but susceptibility to type II, and this contrasting pattern is mainly due to MFOs. This means that enzymes help to detoxify certain types of pyrethroids in some populations but not in others [41,42].

Regarding other enzymes, we observed a significant increase in GST activity in the permethrin-selected HP and LP populations compared with the baseline AF and Rockefeller baseline populations. For lambda-cyhalothrin, no other enzymatic activity than GST was significantly elevated in the HL or LL selected strains compared with the Rockefeller strain, and GST levels were comparable to those observed in the unselected AF strain. GST contributes to insecticide resistance by metabolizing or sequestering insecticides, and elevated levels have been observed in different populations [43]. Lambda-cyhalothrin induces oxidative stress in insects [44], which is countered by enzymes such as MFO and, in particular, GST [45]. The reason the AF population responds differently enzymatically to this insecticide might be that it uses different strategies to overcome oxidative stress that are not detectable by enzymatic assays. Furthermore, although enzymatic assays can provide an overall measure of enzymes such as oxidases, they may not detect specific enzymes. Metabolic resistance is only effective up to the point at which the insecticide exceeds the detoxifying capacity of the enzymes; at higher concentrations, other mechanisms are expected to become dominant.

In the wild, MFOs and GSTs confer pyrethroid resistance across different *Ae. aegypti* populations in Mexico, Brazil [46], Colombia [14], and North America [47]. However, the effects of selective pressure from type II pyrethroids, such as lambda-cyhalothrin, on enzymatic activity require further exploration in strains with diverse genetic backgrounds.

In summary, we showed a tendency for concentration-dependent changes in MFO metabolic activity in the permethrin-selected population, but not in the lambda-cyhalothrin-selected population. Concentration-dependent responses have also been observed in the aphid species *Myzus persicae* [48] when exposed to sublethal concentrations of both type I and type II pyrethroids. This highlights the importance of considering the concentration of insecticides when evaluating metabolic responses. The differing transcription profiles of permethrin and lambda-cyhalothrin suggest that pyrethroids may affect detoxification pathways differently, highlighting the complexity of enzyme activity in these mosquito populations.

### Using transcriptomics to clarify *kdr* and metabolic activity differences

Investigating insecticide resistance mechanisms, including knockdown resistance mutations and enzymatic activity across selected strains, did not explain the higher RR observed in the HP strain compared to the LP strain. Similarly, in lambda-cyhalothrin-pressured strains (HL vs. LL), neither the frequencies of *kdr* alleles and the activity of detoxification enzymes did not correlate well with the observed resistance. Therefore, we used transcriptomics to gain a more comprehensive understanding of these differences. RNA-seq is particularly valuable for detecting differentially expressed genes under diverse stress conditions [23,27,28,49,50] and provides insight into both quantitative and qualitative molecular changes associated with insecticide resistance [51].

Our transcriptome sequencing identified genes whose expression levels exceeded a minimum threshold, thereby conferring insecticide resistance. These genes may contribute to the high level of resistance observed in these phenotypes, which could explain the discrepancies observed.

Insects treated with lambda-cyhalothrin showed a higher number of DEGs (4,103 DEGs) than those treated with permethrin (2,865 DEGs). Despite lambda-cyhalothrin and permethrin sharing a common mode of action [52], the transcriptomic analysis revealed transcriptional differences between the two insecticides.

### Oxidative stress induced by lambda-cyhalothrin activates transcripts that maintain homeostasis and modulate immune pathways

Lambda-cyhalothrin primarily targets the VGSC, but it may also affect chloride and calcium channels [53]. Another molecular target of this insecticide is the respiratory chain complex. Diafenthiuron, propargite, and tetradifon are known to inhibit mitochondrial ATP synthase [54]. Similarly, cyhalothrin, a type II pyrethroid related to lambda-cyhalothrin, has been shown to interfere with the mitochondrial respiratory chain in a concentration-dependent manner [55]. In our study, we identified several genes involved in cellular respiration that were upregulated in response to increasing concentrations of lambda-cyhalothrin. These included cytochrome c oxidase subunit III (AAEL018669) and ATP synthase F0 subunit 8 (AAEL018667).

Several other related genes were also upregulated in response to lambda-cyhalothrin, with the exception of the mitochondrial ATP synthase b chain (AEL005610), which was downregulated. As demonstrated here and elsewhere [15] that lambda-cyhalothrin influences the regulation of transcripts related to the electron transport chain. Additionally, genes associated with the respiratory chain play a role in maintaining cellular homeostasis [56]. A recent study of the plant bug *Lygus pratensis* supports this, finding regulation of oxidative phosphorylation transcripts [57], indicating that our data are not spurious.

Interestingly, reactive oxygen species (ROS) are produced as by-products of oxidative metabolism in response to lambda-cyhalothrin, and their effects have been widely studied in mouse models [44,58]. Once lambda-cyhalothrin enters the cell, it triggers oxidative stress, generating ROS. These reactive oxygen species can damage lipids, DNA, and proteins, inducing apoptosis [44]. In our study, cell death markers, including the caspases (CASPS17, CASPS8, and CASPS19), as well as the initiator caspase Dredd, were found to be upregulated. Additionally, two genes in the NF-κB pathway, were identified: REL1A and REL2. Although they are not significantly overexpressed, they are activated by oxidative damage. These genes, along with serine proteases and clip-domain serine proteases which are abundantly present in our data (S3 Table), are involved in activating the Toll immune pathway. This leads to the production of antimicrobial peptides (AMPs) [59]. We conclude that lambda-cyhalothrin-treated insects may induce oxidative stress that stimulates the Toll pathway component of the innate immune system in unknown ways.

The connections between microbiota and insecticide resistance have recently attracted a great deal of attention. In an unpublished study [60], the authors demonstrate that a seryl-tRNA synthetase-like insect mitochondrial protein paralogue (SLIMP) can regulate the growth of the Bacteroidetes bacterial phylum. Knockdown of SLIMP can promote oxidative stress [61] and is essential for protein synthesis and mitochondrial respiration [62]. Here, we demonstrated that the presence of SLIMP (AAEL006938) is present in lambda-cyhalothrin-resistant strains (S3 Table). Interestingly, a feature of the AF strain used in this study is that antibiotic-induced dysbiosis considerably reduced its resistance status [63]. Resistance could occur if the SLIMP-ROS production modifies bacterial abundances in such a way that only bacterial genera that benefit the insect’s pro-resistance state are present. Future studies could address this hypothesis using RNA interference and quantification of bacterial diversity and abundance instead of antibiotic treatments.

In coordination with the tRNA synthase pathway, we also found that the genes most highly upregulated in lambda-cyhalothrin treatment belonged to transfer RNAs (tRNAs), specifically tRNA-Gly, tRNA-Ser and tRNA-Leu. Notably, these gene transcript levels followed the established pattern. In this sense, tRNA levels correlate with codon usage bias [64], meaning that the tRNA transcript pool could affect protein levels. This is important because more tRNA transcripts could imply faster and more accurate production of proteins that require these specific tRNAs. Although studies on codon usage bias are scarce, permethrin-controlled codon bias of the vgsc channel has been demonstrated in a concentration-dependent manner in *Culex quinquefasciatus* [65]. Codon usage bias has also been associated with sex determination in *Ae. aegypti* [66]. It is worth noting, however, that several serine proteases and clip domain serine proteases were identified in these data, so the presence of tRNA-Ser transcripts is not surprising.

The serine amino acid pathway is equally interesting. Serine is the amino acid that forms the serine and glycine and one-carbon pathway (SGOCP) (reviewed in [67]). In synthesis, the SGOCP pathway converges on many of the resources that insects need. It facilitates the production of nucleotides, proteins, and, interestingly, lipids that form the neurons. The pathway also helps to produce tRNAs in mitochondria (such as SLIMP) and maintains redox balance by regulating NADP and glutathione synthesis. This could explain why the only enzyme that we detected with a significant difference corresponded to GST in lambda-cyhalothrin-treated insects.

In our dataset of genes that are upregulated in response to lambda-cyhalothrin, we identified the sphingomyelin phosphodiesterase gene (AAEL006381 and AAEL009111). This gene is responsible for converting sphingomyelin into its two by-products: phosphocholine and ceramide. Notably, the AAEL006381 gene is upregulated in insects that are refractory to dengue virus infection [68], and it was also strongly associated with pyrethroid resistance in a genetic study [69]. We also observed the downregulation of ceramidase AAEL007030, which was only found in HLR. This could imply that an increase in ceramide levels within cells may have other effects on IR phenotype aside from its role in apoptosis [70] may have other effects on IR phenotype. These genes are highly valuable for functional validation experiments.

In summary, a comprehensive and integrated approach to these pathways is required, particularly with regard to their connection to insecticide resistance, an area that has yet to be explored in insects. This is especially important in the context of oxidative stress induced by lambda-cyhalothrin.

### Potential three-way interaction between OBP-cuticle and CYP4 family transcripts

Following exposure to both insecticides, we observed that many genes related to CYP450 and the cuticle were expressed, which is particularly significant in the context of permethrin resistance. We observed the upregulation of certain CYP450 genes and increased cuticle genes expression at both high and low insecticide concentrations. This is noteworthy because a comparative analysis of *Ae. aegypti* showed that CYP450 genes were upregulated and cuticle genes were downregulated in strains treated with alpha-cypermethrin and lambda-cyhalothrin [27].

This tendency is more pronounced in permethrin-resistant insects. The HPR vs. AF comparison revealed many cuticle genes related to chitin metabolism, as shown by differential expression and GO analyses. A diverse composition of the cuticle is essential for maintaining its hydrophobic properties and plays a key role in communication between external signals and the insect [71]. In this context, genes involved in chitin metabolism that followed this pattern were abundant (Fig 7D and 8C). However, these genes show heterogeneous expression across different studies. AAEL025608 was found to be downregulated in pyrethroid-resistant mosquitoes [72], while another cuticle gene related to AAEL011444 was found to be upregulated in the carcasses of saline-tolerant insects [73]. AAEL022261 was observed to be downregulated in a biopesticide-exposed strain of *Ae. aegypti* [74]. The most highly regulated cuticle proteins, such as AAEL021930 and AAEL025680, were downregulated in *Ae. aegypti* that fed on human blood [75], while AAEL026304 was downregulated in response to flavivirus infection [76]. As no consistent pattern emerges among these genes, we conclude that they are regulated in response to various stressors in these studies.

A growing body of evidence suggests that the thickening or altered structure of the cuticle reduce the penetration of insecticides [71,77,78], thereby promoting insecticide resistance (IR) in *Ae. aegypti*.

Another set of transcripts selected for their relation to IR corresponds to that of Odorant Binding Protein (OBP) transcripts. OBPs are proteins involved in the initial step of chemical perception [79]. They play a role in resistance to xenobiotics in *Tribolium castaneum* [80] and *Spodoptera litura* [81]. OBPs can act as traps that prevent the saturation of neurons and facilitate the transport of molecules [82]. Silencing OBPs in *Cx. quinquefasciatus* results in higher mortality when the insects are treated with deltamethrin [83]. However, in *Ae. aegypti*, OBPs are regulated in various insecticide-selected populations [27]. In our study, we observed the upregulation of OBPs 11, 35, and 21, with OBP 56a being exclusive to HPR. Interestingly, OBPs have been found to potentially bind fatty acids [84]. Furthermore, OBPs bind to insecticides in the cuticle, suggesting an IR mechanism involving OBPs for multiple insecticides [85].

In the context of cuticular metabolism, a family of CYP450 enzymes has been implicated in the production of cuticular hydrocarbons in *An. gambiae* [86]. The CYP enzyme specifically involved in this process belongs to family 4, CYP4G16. We found that the expression of genes in the CYP4 family was increased in all permethrin-treated insects compared with the AF strain. Notably, only survivors of the high-resistance HPR strain exhibited upregulation of CYP4G36. Another gene, corresponding to CYP4H32, maintained the HPR > HPS > LP pattern. Few studies have characterized the cuticle and its composition in relation to resistance mechanisms.

Finally, our results suggest that a multigenic response to pyrethroid insecticides occurs in *Aedes aegypti*, particularly at high insecticide concentrations. At low concentrations, mechanisms such as *kdr* and metabolic resistance are relevant, but at high concentrations, other metabolic pathways are activated (Fig 9). Further research is needed to explore the interactions between cuticle proteins, OBPs, and CYP450s in response to insecticides. This is particularly important for strains whose resistance mechanisms are not well understood and deviate from classical mechanisms such as *kdr* and metabolic resistance. This research will help to establish a relationship with the resistant phenotype in *Ae. aegypti*.

**Fig. 9.**
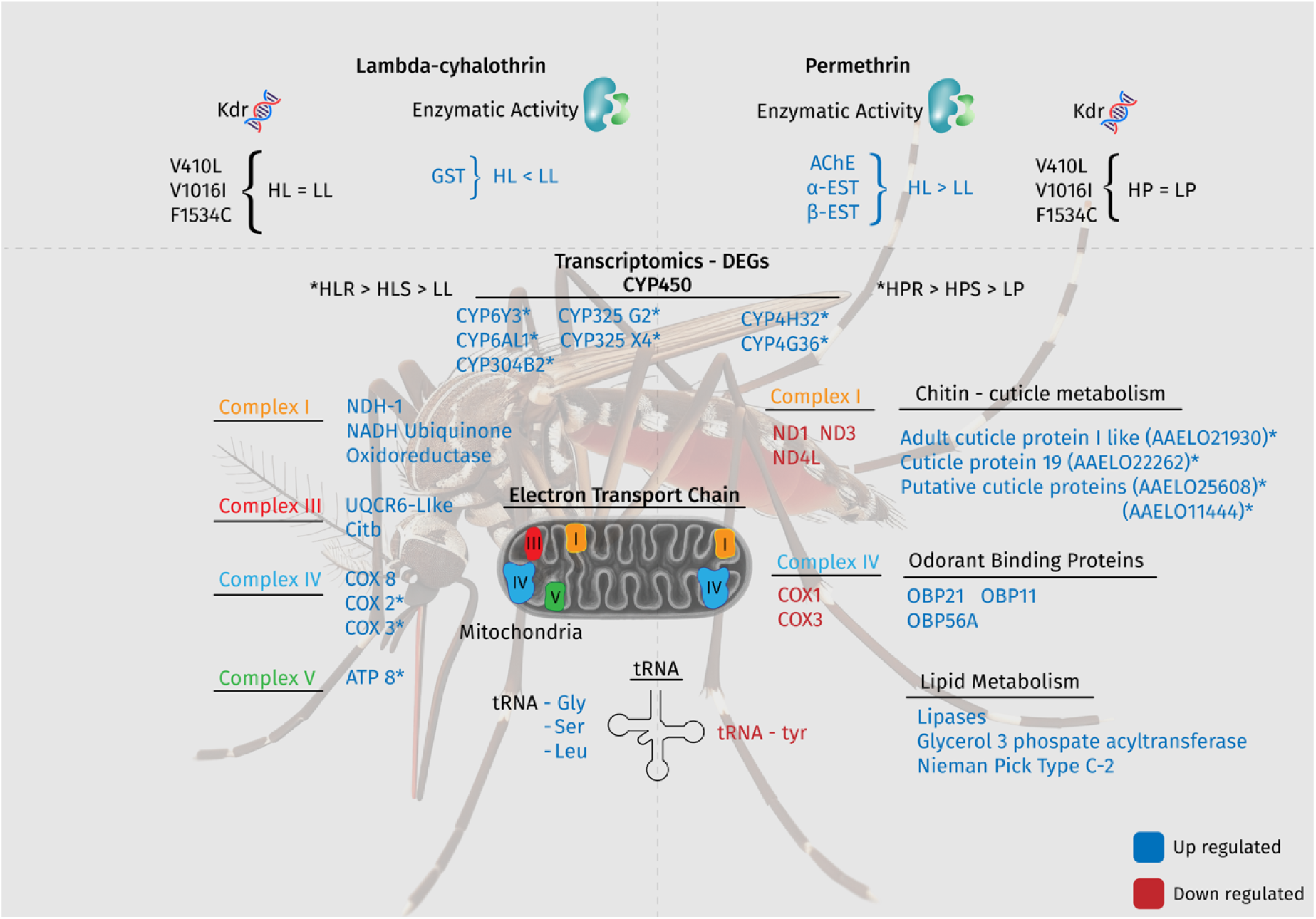
Summary of the differentially expressed transcripts in *Aedes aegypti* mosquitoes exposed to the insecticides lambda-cyhalothrin (left) and permethrin (right). The frequency of the *kdr* allele in highly resistant mosquitoes (HL and HP) is consistent with that in susceptible mosquitoes (LL and LP).

## Conclusions

By analysing these *Ae. aegypti* strains at different insecticide concentrations, we have identified transcripts that respond sequentially and which may be required at low levels to overcome insecticides. These findings provide insight into the mechanistic links between established and emerging insecticide resistance mechanisms, including oxidative damage and its connection to immune pathways triggered by insecticides. Further research into gene–transcript–protein interactions is needed to gain a deeper understanding of the insecticide resistance phenotype. Additionally, it is crucial to examine regulatory modifications within these pathways, since transcriptomic data only offer a snapshot of expression patterns. A more thorough investigation of the underlying regulatory networks will provide valuable insights into the complexity of these traits.

These results also emphasize the importance of public health agencies implementing more thorough, consistent and widespread insecticide spraying strategies. Our experiments demonstrate that *Ae. aegypti* responses vary according to the type of pyrethroid and the concentration of the insecticide, emphasising the need for optimized protocols to effectively control resistance.

## Notes

### Competing Interest Statement

The authors have declared no competing interest.

